# Ebselen attenuates mycobacterial virulence through inhibition of ESX-1 secretion

**DOI:** 10.1101/828574

**Authors:** Morwan M. Osman, Malte L. Pinckert, Sew Peak-Chew, Mark A. Troll, William H. Conrad, Lalita Ramakrishnan

**Affiliations:** Molecular Immunity Unit, Department of Medicine, University of Cambridge, MRC Laboratory of Molecular Biology, Cambridge CB2 0QH, UK; MRC Laboratory of Molecular Biology, Francis Crick Avenue, Cambridge CB2 0QH, UK; York Structural Biology Laboratory, Department of Chemistry, University of York, York, YO10 5DD, UK; Division of Virology, Department of Pathology, Addenbrooke’s Hospital, University of Cambridge, Cambridge, CB2 0QQ, UK; Department of Chemistry, Lake Forest College, Lake Forest, Illinois 60045, US

## Abstract

The type VII secretion system ESX-1 mediates virulence in *Mycobacterium tuberculosis* and *Mycobacterium marinum*. We find that in *M. marinum*, the synthetic organoselenium compound ebselen inhibits secretion of ESAT-6, a major ESX-1 substrate. We find that ebselen inhibits the *in vitro* activity of the ESX-1 AAA+ ATPase EccA1, which potentiates ESX-1 substrate secretion and function. Ebselen modifies a cysteine in its N-terminal tetratricopeptide repeat domain that is required for EccA1’s *in vitro* ATPase activity. Surprisingly, mutational analyses show this this cysteine is not required for ESX-1 secretion or ebselen’s activity, showing that ebselen inhibits ESX-1 secretion independently of inhibiting EccA1 activity *in vitro*. While the mechanism by which ebselen inhibits ESX-1 secretion remains elusive, we show that it attenuates ESX-1-mediated damage of *M. marinum*-containing macrophage phagosomes and inhibits intramacrophage growth. Extending our studies to *M. tuberculosis*, we find that ebselen inhibits ESX-1 secretion and phagosomal membrane damage in this organism. This work provides insight into EccA1 biology. Ebselen is an orally active drug in clinical trials for other conditions and this work suggests its potential in tuberculosis therapy.

## INTRODUCTION

*Mycobacterium tuberculosis* (Mtb) and its close genetic relative *Mycobacterium marinum* (Mm) require their type VII secretion system ESX-1 (ESAT-6 Secretion System 1) for virulence[1–3]. ESX-1 was first identified as a virulence determinant when it was shown that a 9.4 kb deletion (ΔRD1) in its locus was the primary cause of attenuation for the live attenuated vaccine BCG [1,3–6]. ESX-1 mediates membranolytic activity reflected by damage to the membranes of mycobacterium-containing macrophage phagosomes[7,8]. This damage is thought integral to ESX-1 mediated virulence functions such as intramacrophage growth[9]. *In vitro*, ESX-1 mediates membrane disruption of infected lung epithelial cells [10], cultured macrophages [10], and red blood cells (RBCs) [11–13]. Previously, ESX-1 membranolytic activity had been ascribed to its secreted substrate ESAT-6 forming pores in host membranes[10,12,14]. In 2017, we found that the pore-forming activity ascribed to ESAT-6 was due to residual detergent contamination of ESAT-6 preparations[15]. Moreover, we found that ESX-1 membrane disruption was exclusively contact-dependent and caused gross membrane disruptions as opposed to distinct pores [15].

In this paper, we report studies that were instigated by our speculation that ESX-1 might mediate membrane disruptions through peroxidation of host membrane lipids. Following this hypothesis, we identified the antioxidant drug ebselen (2-phenyl-1,2-benzisoselenazol-3(2H)- one) as an inhibitor of *M. marinum* (Mm) ESX-1’s’s hemolytic function. We determined that ebselen inhibition of ESX-1 substrates, including ESAT-6. In searching for the relevant target of ebselen, we investigated the ESX-1 ATPase EccA1, and found that it specifically inhibited its *in vitro* ATPase activity through covalent binding to a cysteine in its proposed substrate binding domain. However, ebselen’s effect on ESX-1 secretion in Mm was not due to EccA1 inhibition. Finally, we showed that ebselen inhibits ESX-1 function within infected macrophages, with treatment inhibiting phagosomal damage and intramacrophage growth. Finally, we found that ebselen also inhibits ESAT-6 secretion and phagosomal damage in Mtb, suggesting its potential as an adjunctive treatment for tuberculosis.

## RESULTS

### Ebselen inhibits ESX-1 secretion in *Mm*

To test the hypothesis that ESX-1-dependent membrane disruptions could occur through peroxidation of host lipids, we screened antioxidants for their ability to inhibit Mm’s contact-dependent hemolytic activity. We tested the antioxidants ebselen, ascorbate, and butylated hydroxytoluene, and only ebselen inhibited Mm’s RBC lysis with a half maximal inhibitory concentration (IC_50_) of 9.3 μM (Fig. 1A). These findings indicated that ebselen’s effects on RBC lysis was independent of its antioxidant effect, suggesting that lipid peroxidation was not the mechanism behind ESX-1 membrane disruption.

**Figure 1:**
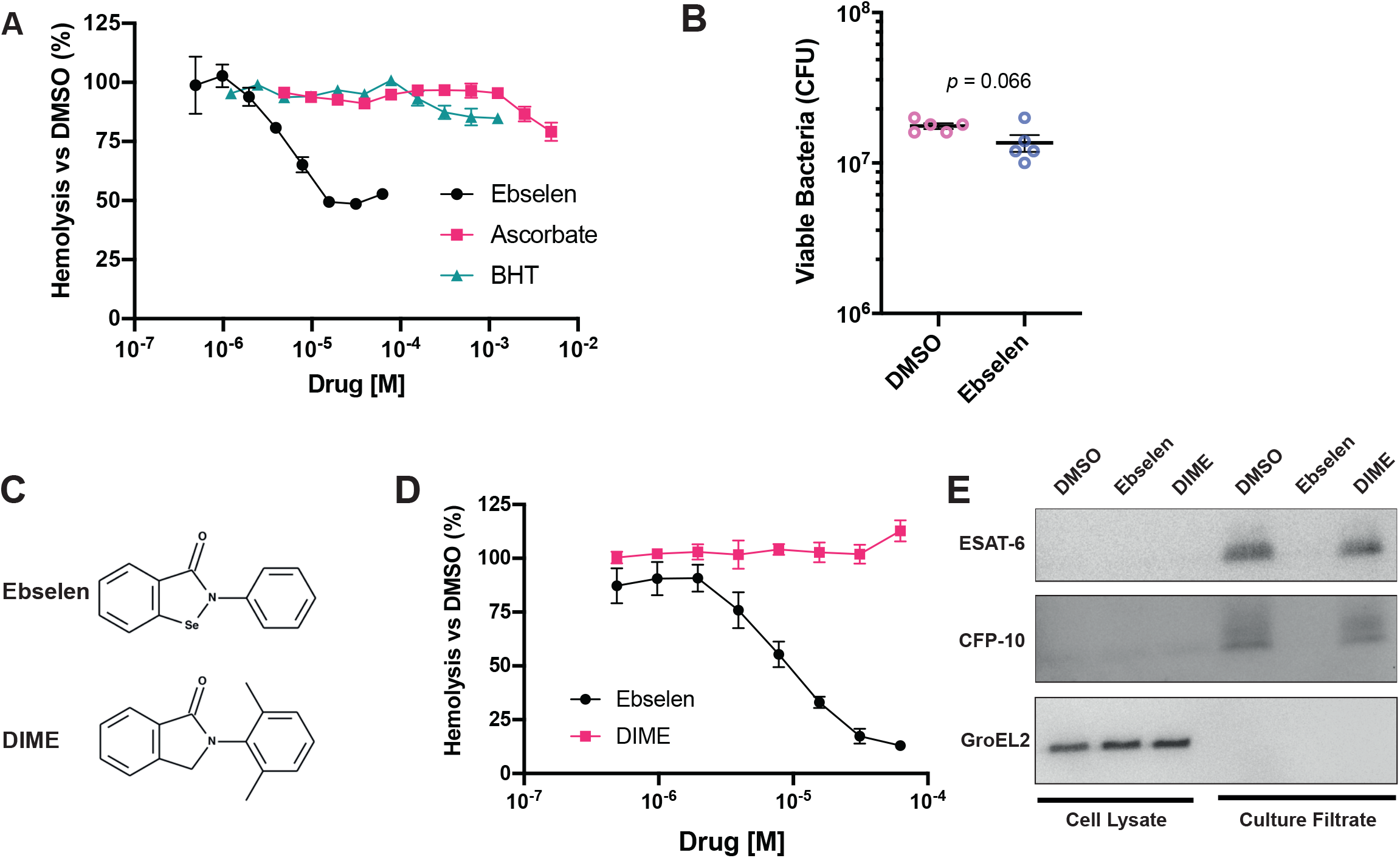
Ebselen treatment inhibits ESX-1 Mediated Membrane Disruption & Secretion. **A:** Percent sRBC hemolysis by *M. marinum* relative to DMSO treatment in the presence of increasing concentrations of antioxidants. Data are presented in biological triplicate. **B:** Structures of ebselen and analogue 2-(2,6-dimethylphenyl)-1-isoindolinone (DIME) **C:** Colony forming units of *M. marinum* following 4 hours of exposure to 62.5 μM Ebselen or 0.5% DMSO. Data representative of 3 independent experiments. Student’s t-test. **D:** Percent maximum hemolysis by wildtype *M. marinum* in response to ebselen or DIME treatment. Data presented in biological triplicate **E:** Western blot of lysed *M. marinum* cell pellet and culture filtrate fractions after 4 hours incubation with 32 μM ebselen, DIME, or 0.5 %DMSO. anti-ESAT-6; anti-CFP-10 and anti-GroEL2 (loading control). Blot is representative of 3 independent experiments. All error bars represent SEM.

We pursued how ebselen might inhibit hemolysis independent of its antioxidant effect. As ebselen has been reported to have antimicrobial activities against Mtb [16], we set to determine if bacterial killing was responsible for ebselen’s effect on Mm’s hemolytic activity. We plated Mm that had been exposed to ebselen for the duration of the hemolysis assay and found that even the highest concentration used, 62.5 μM, did not affect viability (Fig. 1A and 1B). Ebselen’s drug-like activities are mediated through its selenium moiety, which enables it to covalently modify free thiols, such as free cysteine residues [17,18]. Consistent with this, we found that an ebselen analog lacking the selenium moiety, 2-(2,6-dimethylphenyl)-1-isoindolinone (DIME) failed to inhibit hemolysis (Fig. 1C and D). We next asked if ebselen’s primary action was on ESX-1 secretion, assessing the secretion of two co-secreted substrates, ESAT-6 and CFP-10, as a proxy for overall ESX-1 secretion. Ebselen treatment inhibited both ESAT-6 and CFP-10 secretion, while DIME had no effect (Fig. 1E). Altogether, these results show that ebselen inhibits ESX-1 secretion likely by cysteine modification.

### Ebselen inhibits EccA1’s *in vitro* ATPase activity

Ebselen has been shown to inhibit the activity of prokaryotic and eukaryotic enzymes, including the yeast plasma membrane H+-ATPase [19]. These inhibitory functions occur through ebselen’s reactivity with free thiols, including free cysteine residues [18]. ESX-1-mediated secretion is dependent on two ATPases within the locus: the FtsK/SpoIIIE ATPase EccC1, encoded by the adjacent genes EccCa1 and EccCb1, which powers the transport of ESX-1 substrates[20–22], and the AAA+ ATPase EccA1, which is thought to act as a chaperone for ESX-1 substrates [21,23,24]. The ATPase activity of purified Mtb EccA1 protein has been demonstrated [25]. We attempted to purify both EccA1 and EccC1 (as an EccCa1/Cb1 fusion) and only succeeded in purifying EccA1. We found that ebselen inhibited the ATPase activity of purified EccA1 in a dose-dependent manner, with an IC_50_ of 2.5 μM (Fig. 2A). To determine if this inhibition was via cysteine modification, mass spectrometry analyses were performed on the samples treated with the various ebselen concentrations in Fig. 2A. We identified ebselen-modified peptides corresponding to cysteines 204 and 531 (Figure 2B, S1).

**Figure 2:**
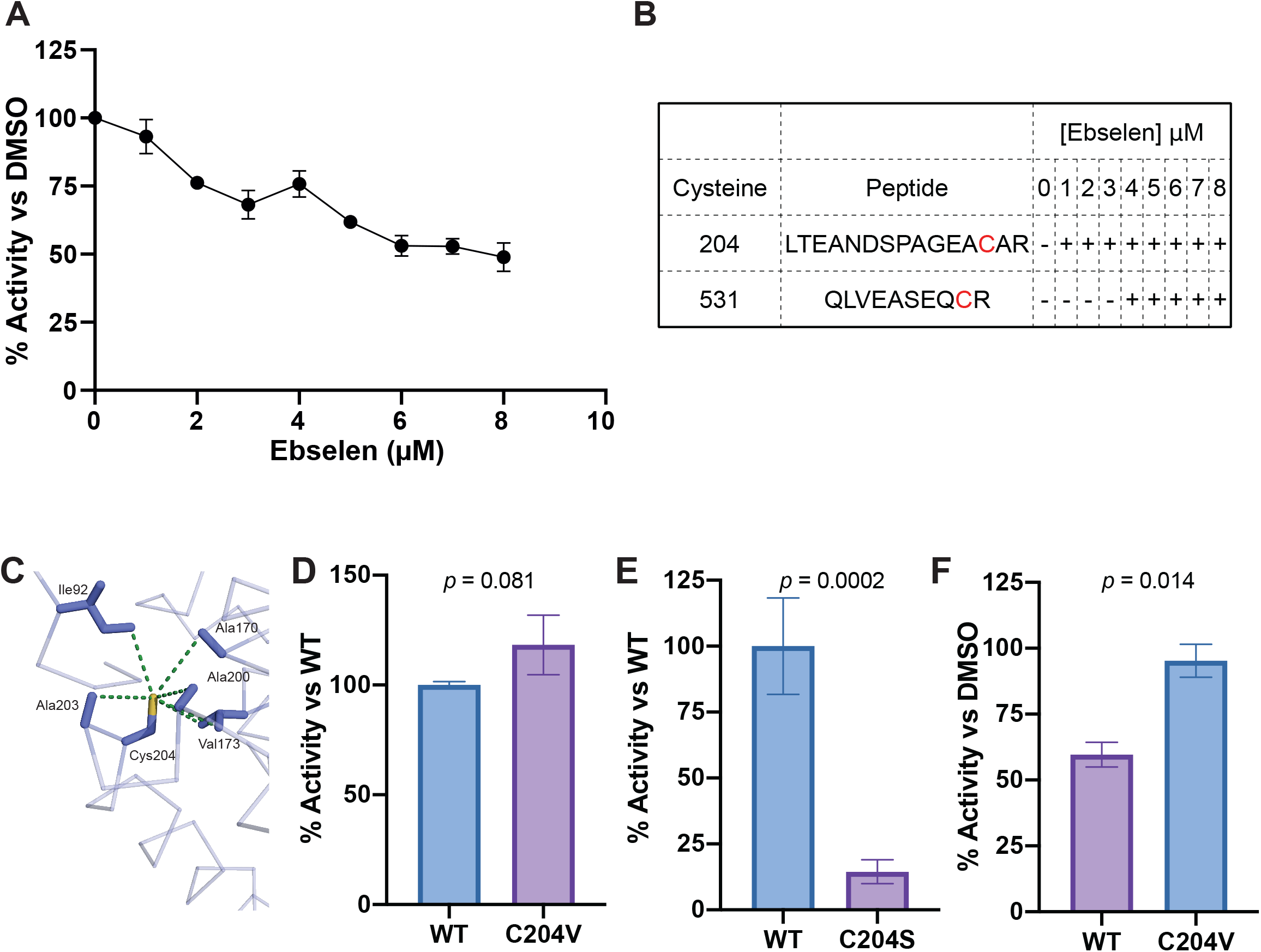
Ebselen inhibits EccA1 through covalent modification of Cys204 and Cys531. **A:** Dose-response curve of EccA1 treated with ebselen. ATPase activity is normalized to EccA1 treated with DMSO. Data are biological triplicate. Error bars show SEM. **B**: Qualitative representation of ebselen modified peptides containing Cys204 and Cys531 detected by MS at increasing concentrations of ebselen. + symbols indicate concentrations at which ebselen modified peptides were observed. **C**: Model of Cys204 of EccA1 (PDB accession code 4F3V) and residues located within 5 angstroms, with dashes measuring distance. **D:** ATPase activity of EccA1 C204V point mutant (C204V) vs wildtype EccA1 (WT). Activity normalized to WT. Data from two independent experiments. ATPase activity is normalized to wildtype. Student’s t-test. **E**: ATPase activity of EccA1 C204S point mutant (C204S) compared to wildtype EccA1 (WT). ATPase activity is normalized to wildtype. Data from five independent experiments. Student’s t-test. **F:** Relative ATPase activity of EccA1 C204V and wildtype EccA1 (WT) treated with 8 μM ebselen. ATPase activity is normalized to DMSO treatment for each protein. Data from three independent experiments. Student’s t-test. Error bars show SEM, p values are indicated.

In addition to the ATPase domain that mediates EccA1’s assembly and enzymatic activity [25], EccA1 has a large N-terminal tetratricopeptide repeat (TPR) domain predicted to mediate substrate binding [26] and is dispensable for its *in vitro* ATPase activity[25]. Of the two residues modified by ebselen, only Cys204 is present in Mm (Figure S2), and we found that ebselen’s binding to Cys204 was also linked to inhibition of activity, suggesting that conservation of the TPR domain is required for EccA1’s ATPase activity. This was supported by the MS data indicated that ebselen modification of Cys204 preceded that of Cys531 (Figure 2B, S1).

Examination of the crystal structure of EccA1’s TPR domain (PDB: 4F3V) revealed that Cys204 is located in the TPR domain’s hydrophobic core (Fig. 2C) [26]. To confirm Cys204’s role in ebselen’s activity, we engineered EccA1 where the Cys204 had been changed to a valine (EccA1 C204V) or to the more hydrophilic residue serine (EccA1 C204S). We saw that EccA1 C204V had similar ATPase activity to the wildtype enzyme, while EccA1 C204S showed only 14% of wildtype EccA1 (Figure 2D,E). These findings suggested that while Cys204 is dispensable for enzymatic activity, conservation of its hydrophobic environment is essential for EccA1’s *in vitro* ATPase function.

If ebselen’s ATPase inhibitory activity was primarily due to modification of Cys204, then EccA1 C204V should not be inhibited by ebselen treatment. When we treated both wildtype and EccA1 C204V with 8 μM ebselen, we found that EccA1 Cys204 was essential for ebselen inhibition: with EccA1 C204V retaining 95% of its ATPase activity vs ∼60% for WT EccA1 (Figure 2F). We expected to see an intermediate level of inhibition, as we identified Cys531 modified peptides in wildtype EccA1 treated with 8 μM ebselen. Instead, EccA1 C204V’s insensitivity indicates that Cys204 modification must either precede Cys531 modification or that Cys531 modification does not contribute substantially to ebselen’s inhibitory effect. Together, these results show EccA1 Cys204 is essential for ebselen’s *in vitro* inhibition of EccA1 ATPase activity.

### Ebselen’s effect on ESX-1 secretion is independent of EccA1

We next asked whether the loss of *in vitro* ATPase activity was the mechanism for ebselen’s inhibition of ESX-1 secretion. We generated an unmarked Mm-Δ*eccA1* mutant by excising the hygromycin cassette from an Mm-*eccA1*::Tn mutant, and confirmed it was still deficient in secretion of ESAT-6 and CFP-10 (Figure 3A). We then complemented Mm-Δ*eccA1* with the plasmids pMOFXh-*eccA1*_Mtb_ and pMOFXh-*eccA1*_Mtb_-C204V, which express Mtb EccA1 and EccA1-C204V respectively. Mm-::EccA1_Mtb_ and Mm-::C204V both restored secretion of ESAT-6 (Fig 3B). If ebselen’s ESX-1 inhibitory effects are mediated through EccA1, then Mm expressing EccA1-C204V should secrete ESAT-6 in the presence of ebselen. To our surprise, we found that ebselen treatment inhibited ESAT-6 secretion similarly in Mm-Δ*eccA1*::C204V as in wildtype Mm and Mm-Δ*eccA1*:: EccA1_Mtb_. Thus, ebselen’s effects on ESX-1 secretion were not through inhibition of EccA1. Furthermore, complementation Mm-Δ*eccA1* with EccA1-C204S rescued secretion, demonstrating that the *in vitro* ATPase defects we observed were not sufficient to abrogate EccA1’s role in ESX-1 function.

**Figure 3:**
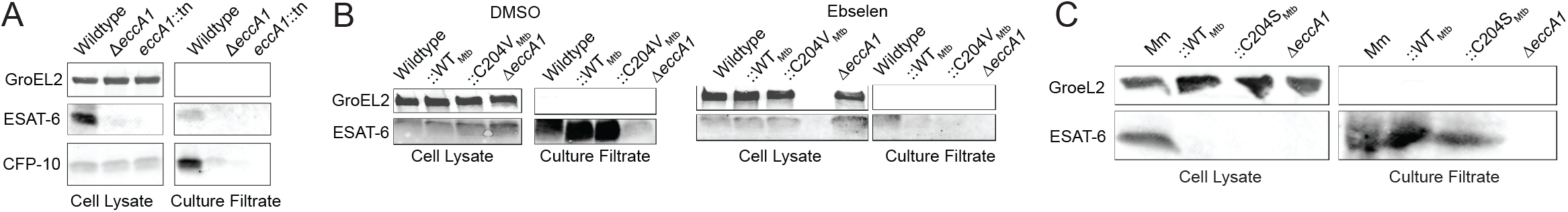
EccA1’s role in ESAT-6 secretion does not correlate with its in vitro ATPase activity. **A:** Immunoblots of Mm cell lysate (left) and culture filtrate (right) fractions. anti-GroEL2 (loading control). **B:** Fractions of Mm treated with 0.5% DMSO or 32 μM ebselen for 4 hours. **C**: Fractions of Mm & Mm mutants after 48hours of growth in Sauton’s Medium. GroEL2 is used as a loading control.

### Ebselen inhibits ESX-1-mediated virulence phenotypes in Mm-infected macrophages

Our inability to find ebselen’s target notwithstanding, its inhibitory effects on ESX-1 secretion suggested it could be a promising antitubercular drug. We asked if ebselen treatment of Mm infected macrophages would inhibit ESX-1 virulence-associated functions in infected macrophages. We tested its effects on phagosomal damage of mycobacterial compartments, thought to be primary to ESX-1-dependent intramacrophage growth[9]. We used the galectin 8 assay to determine whether ebselen treatment affected the extent to which Mm could damage its resident compartment within the macrophage. Galectin 8 is recruited to sites of membrane damage and is a sensitive measure of phagosomal damage in Mm- and Mtb-infected macrophages[27,28]. We found that treatment with 16 μM ebselen significantly reduced the amount of phagosomal damage induced by wild-type Mm (Fig 4A), demonstrating that ebselen effectively inhibits Mm ESX-1 in the context of the infected macrophage. Moreover, ebselen also inhibited intramacrophage growth (Figure 4B). However, this reduction, though significant. was partial, being much less than that of the Mm-ΔRD1 strain (45% vs. 92% reduction over 3 days), and moreover, there was no significant difference in growth for concentrations ranging from 16 to 64 μM, showing that we had identified its maximal effect in this assay (Fig 4B).

**Figure 4:**
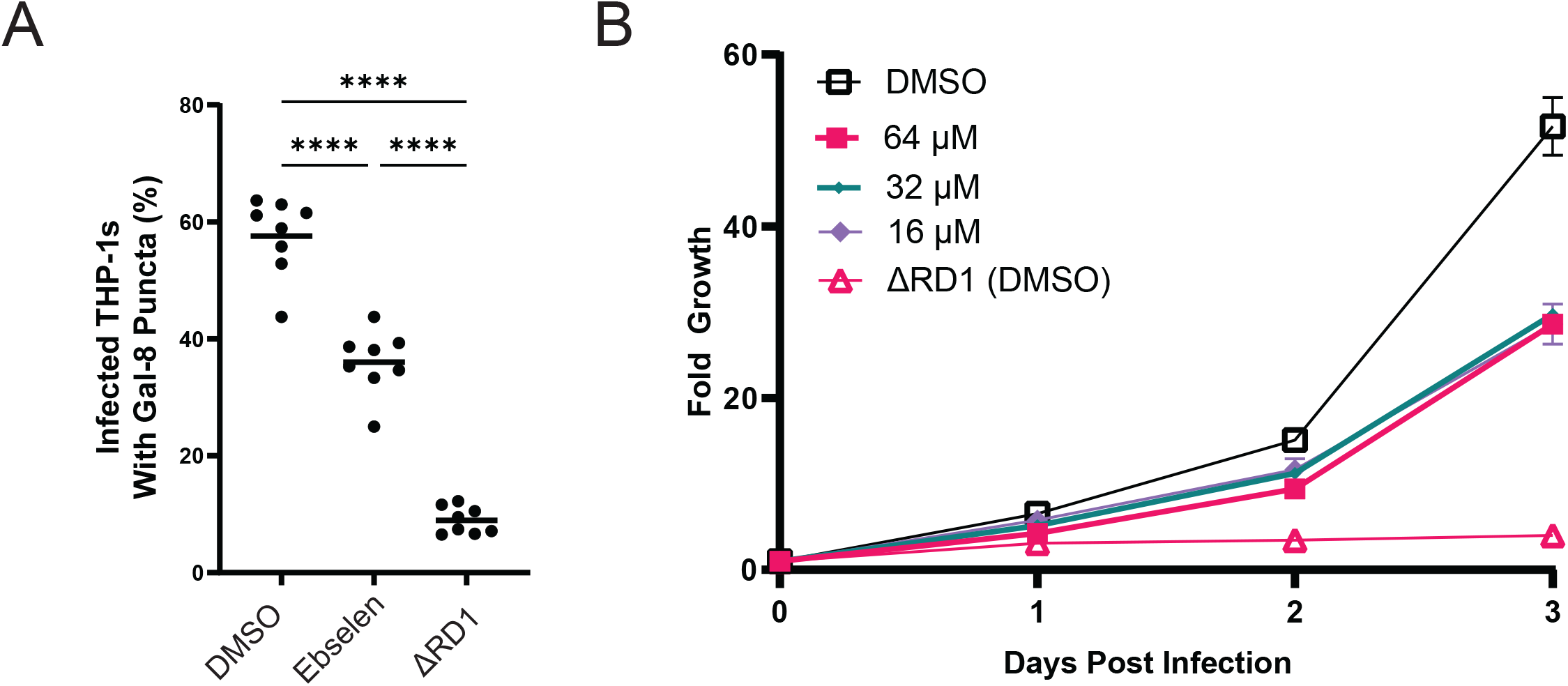
Ebselen inhibits Mm ESX-1 function in macrophages. **A:** % of Mm-infected THP-1 macrophages with galectin-8 puncta. DMSO – wildtype Mm treated with 0.5% DMSO. Ebselen – 16 μM Ebselen. Data representative of three independent experiments. **B: :** Intramacrophage growth of ebselen-treated Mm within J774A.1 cells as measured by bacterial fluorescence. Doses shown in legend. Data representative of three independent experiments.

### Ebselen inhibits Mtb ESX-1 function

We have previously shown that *Mm* and *Mtb* ESX-1 are functionally equivalent, with the Mtb ESX-1 locus capable of complementing an ESX-1-deficient Mm RD1 deletion (Mm-ΔRD1) mutant’s defects in hemolysis, secretion, and virulence [15]. To determine if ebselen inhibited ESX-1 secretion in *Mtb*, we used the Mtb H37Rv derivative mc^2^6206, which is auxotrophic for leucine and pantothenic acid but retains an intact ESX-1 locus [29]. The mean inhibitory concentration (MIC) of ebselen has been measured to be significantly lower in Mtb, with values ranging from 36.4 to 76.4 μM depending on strain [30,31] versus 100 μM for Mm, we selected doses of 8 and 16 μM to ensure any effects we were seeing were not due to the microbicidal activity of ebselen. We found that at both doses, ebselen treatment reduced ESAT-6 secretion (Fig 5A). We then set out to determine whether these doses were capable of inhibiting Mtb’s function during macrophage infection. We turned to the galectin 8 assay to measure the extent that ebselen treatment could inhibit ESX-1 function within the macrophage. We found that Mtb, both doses significantly reduced the frequency of phagosomal damage by Mtb, with 16 μM showing a greater reduction than 8 μM (28.4% vs 36.5% of infected cells with puncta) (Figure 5B). Together, these results demonstrate that ebselen effectively inhibits the ESX-1 function of Mtb.

**Figure 5:**
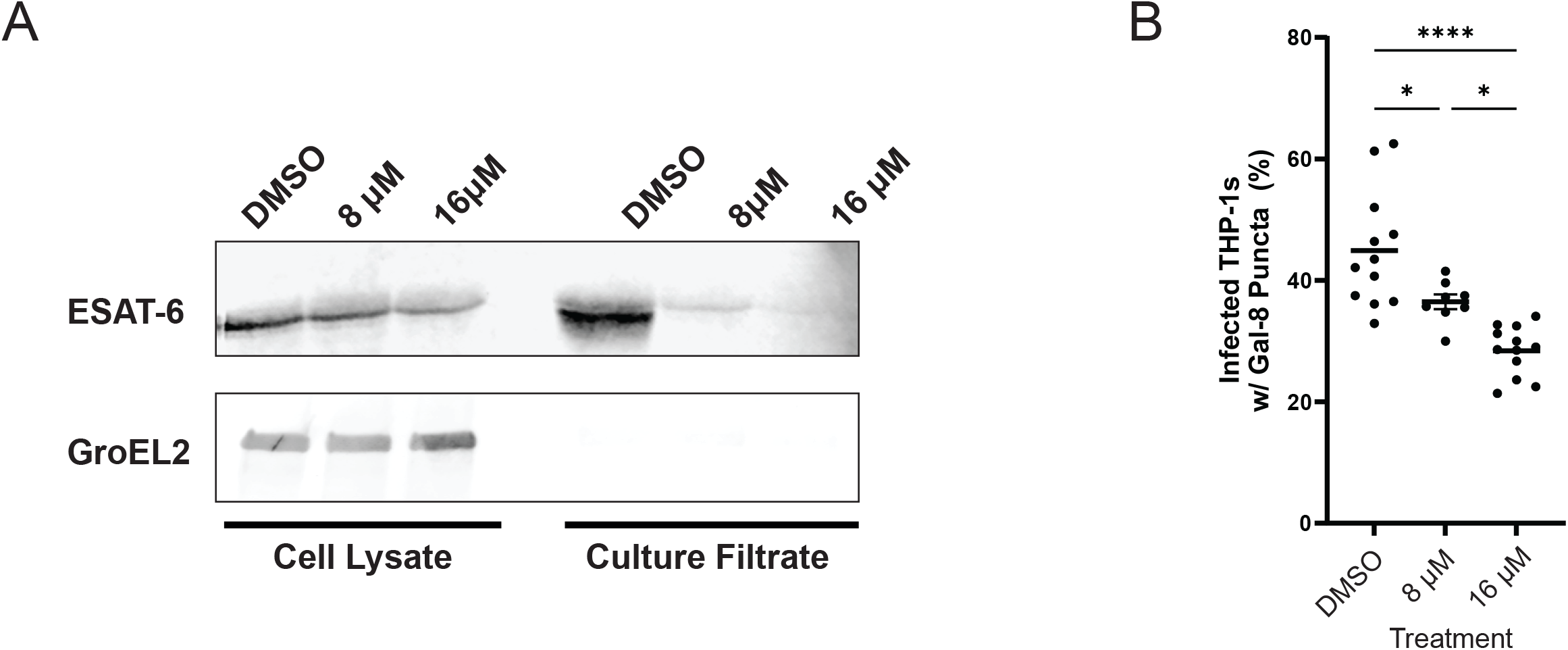
Ebselen inhibits ESX-1 function in *M. tuberculosis*. **A:** Western blots of lysed *M. tuberculosis* cell pellet and culture filtrate fractions after 96 hours. Blot representative of three independent experiments. **B:** % of Mtb-infected THP-1 macrophages with galectin-8 puncta. Labels correspond to treatment with 0.5% DMSO, 8 μM Ebselen, or 16 μM Ebselen. Data representative of three independent experiments.

## Discussion

This work identifies ebselen, a drug in clinical trials for many different medical conditions (NCT:04677972, 02819856,03013400, 05117710), as a potential adjunctive treatment for TB. We showed that ebselen works by inhibiting ESX-1 secretion and function. Because the ESX-1 secretion system is a critical virulence determinant, there has been a systematic study to screen FDA-approved drugs that inhibit Mtb ESX-1 secretion [32]. This screen found that ethoxzolamide (ETZ), an FDA-approved drug, inhibits ESX-1 secretion via suppression of PhoPR signaling [32]. However, we have found that PhoPR does not regulate ESAT-6 secretion in Mm as it does in Mtb (F. Chu, C. Cosma and LR, unpublished data). Because ebselen inhibits ESX-1 secretion in both Mm and Mtb, we can deduce its mechanism is distinct from ETZ’s. Rybniker et. al. identified ESX-1 inhibitors that operate through different mechanisms: BTP15 and BBH7. BTP15 which inhibits the MprB histidine kinase which regulates the ESX-1 *espACD* operon, while BBH7 disrupts mycobacterial metal homeostasis, resulting in dysregulation of ESX-1 secretion [33]. Our discovery of ebselen, in contrast, was serendipitous, resulting from searching for the mechanism of mycobacterial ESX-1-mediated membrane disruption. Moreover, despite our efforts, we have not yet identified ebselen’s target. Our initial data suggesting ESX-1 AAA+ ATPase EccA1 as its target turned out to be a false lead. However, this finding did shed light on EccA1 biology and suggests that either its residual activity with ebselen treatment (∼14%) is sufficient for secretion, and/or that its activity, like that of other AAA+ ATPases, is induced by specific *in vivo* substrates missing in our *in vitro* biochemical activity assay[34]. Despite not knowing its target, our finding that this safe, well-tolerated drug[35,36] can work as an anti-virulence drug in TB is potentially important. Ebselen has promiscuous activity, reacting with any free sulfhydryl group and therefore can inhibit multiple human enzymes including inositol monophosphatase [37] and indoleamine 2,3-dioxygenase [38]. This may explain its promise in multiple conditions ranging from Meniere’s disease, drug-induced ototoxicity, bipolar disorder and treatment-resistant depression (NCT: 04677972, 02819856, 03013400, 05117710).

Ebselen has been demonstrated to show antimicrobial activity in *Helicobacter pylori* and *Mtb* [39]. Its antimicrobial effects on Mtb have been attributed to covalent modification of the essential Antigen 85 complex (Ag85C) [30]. Ag85C synthesizes the lipid trehalose dimycolate (TDM) from trehalose monomycolate, and ebselen treatment abrogates this activity [30]. However, ebselen’s inhibition of Mtb TDM synthesis is observed only at concentrations at or above 73 μM [30], whereas we observed inhibition of Mtb ESX-1 secretion at 8 μM concentrations. Thus, its inhibition of ESX-1 secretion is distinct from its inhibition of Ag85C.

Ebselen has been demonstrated to be clinically safe and well tolerated in humans [35], suggesting its potential to standard anti-tuberculous chemotherapy. In Meniere’s disease which can be associated with hearing loss, a recently completed phase 2B trial found that ebselen achieved pre-specified end points in improving this hearing loss relative placebo (NCT:04677972). Ebselen has been shown to be otoprotective against aminoglycoside-induced ototoxicity in mice through its antioxidant activity[40]. This finding is being followed up in an ongoing clinical trial to see if it prevents aminoglycoside-induced ototoxicity in cystic fibrosis patients (NCT: 02819856). Aminoglycoside-induced ototoxicity is difficult to predict and presents a challenge in completing aminoglycosides-containing regimens, the mainstay of drug-resistant TB treatment [41]. Our findings that it may suggests its otoprotective, antimicrobial, and anti-virulence properties may combine to make it particularly attractive in aminoglycoside containing tuberculosis treatment regimens.

## Materials and Methods

### Bacterial strains and methods

Strains used are listed in Supplementary Table 2. All Mm strains were derived from wild-type Mm purchased from American Type Culture Collection (strain M, ATCC #BAA-535). Wildtype Mm was maintained as described previously[15]. Mm-*eccA1::*Tn was pulled from an Mm transposon mutant library (C. L. Cosma, L.R., unpublished data). Mm- Δ*eccA1* was generated by excision of the mariner-transposon via transformation with pYUB870. Loss of pYUB870 was confirmed by plating on 7H10+sucrose plates, and loss of the transposon was confirmed via PCR and loss of growth on selective media. *M. tuberculosis* mc^2^6206 was cultured in 7H9 complete media containing 0.05% tween 80 supplemented with pantothenic acid (12 μg/mL) and leucine (25 μg /mL).

### Hemolysis assay

Hemolytic activity was assessed as described previously[15]. Briefly, 100 μL of sheep red blood cells (sRBCs) were transferred to each condition and 100 μL of PBS or bacterial suspension were added on top and incubated for two hours at 33°C. 100% lysis with 0.1% Tx100 (Sigma). Drugs stocks were made at 200x concentrations and 0.5% DMSO was used as a vehicle control.

### Secretion Assays

Secretion assays were conducted as described previously with minor modifications [15]. Briefly, Mtb mc^2^6206 and Mm were grown to mid to late log stage and washed with PBS before being resuspended to a final OD_600_ of 0.8. *M. tuberculosis* mc^2^6206 was resuspended in 50 mL 7H9 media supplemented with 10% DC (2% dextrose, 145 mM NaCl, 30 μg/mL bovine catalase), pantothenic acid, (12 μg/mL) and leucine (25 μg /mL). Cultures were incubated for 4 days at 37°C in the presence of drug or 0.5% DMSO. Mm was resuspended in 50 mL Sauton’s Media and incubated for 4 hours at 33°C in the presence of drug or 0.5% DMSO. Culture filtrate (CF) and cell pellet (CP) fractions were prepared as described previously [15]. 10 μg of CP and 30 μg of CF were loaded per well for SDS-PAGE, and presence of ESAT-6, CFP-10 and GroEL2 were determined by western blotting with mouse anti-ESAT-6 clone 11G4 (1:1,000; Thermo Fisher, HYB-076-08-02), rabbit anti-CFP-10 (1:500; BEI, product NR13801), or mouse anti-GroEL2 clone IT-56 (1:1,000; BEI, product NR-13655).

### Purification of EccA1

Purification constructs were expressed in *E. coli* C43 (DE3) (Lucigen). Single colonies were used to inoculate 100 mL of Lysogeny Broth (LB) media containing 50 μg/mL Kanamycin and incubated overnight (O/N) at 37°C. 1L of Terrific Broth (TB) was inoculated at an OD_600_ of 0.015 and grown at 37°C to an OD_600_ between 0.6-0.8 and cooled to 20°C and induced with 1 mM IPTG O/N. Pelleted cells were resuspended in ice-cold Lysis Buffer (2.5xPBS; 10 mM Imidazole) with SIGMAFAST™ Protease Inhibitor tablets (Sigma). Cells were lysed in three passages through an Avestin Emulsiflex C3 at 15,000 psi. Debris and unbroken cells were removed by centrifuging at 18,000 x g for 20m. ATP and MgCl_2_ were added to lysate to a final concentration of 5 and 20 mM respectively, and lysate was bound to a HisTrap HP column (GE Life Sciences). Column was washed 3x with 10 Column Volumes (CV) of Wash Buffer [2.5 x PBS; 40 mM Imidazole] and eluted with 12 CV of Elution Buffer [2.5x PBS; 500 mM Imidazole]. Eluted fractions were concentrated to 2 mL with 30 kDa cutoff concentrators (Amicon) and diluted 1:10 in IEX1 buffer [20mM Tris pH 8.0 (RT) ; 15mM NaCl]. ATP and MgCl_2_ were then added to a final concentration of 1 and 20 mM and then applied to a HiTrap Q column (GE Life Sciences). Separation occurred over an 18 CV gradient from 100% IEX1 to 100% IEX2 buffers [20mM Tris pH 8.0 (RT) ; 500mM NaCl]. Peak fractions were concentrated with 30 kDa cutoff concentrators (Amicon) to ∼10 mg/ml and loaded onto a S200 Increase 10/300 GL (GE Life Sciences) in SEC Buffer [20mM Tris, pH 8.0 (RT); 100mM NaCl]. Peak fractions were pooled and concentrated, and aliquots were flash frozen and stored at -70°C.

### ATPase Assays

ATPase activity of EccA1 and EccA1 point mutants was measured with an ATPase/GTPase kit (Sigma) adapted for 384 well-plates. Experiments were conducted for 180 minutes with a final EccA1 concentration of 1 μM. For ebselen treatments, samples were incubated on ice with drug for 20 min prior to assay. Background activity was determined by conducting reactions in the absence of added ATP.

### Mass Spectrometry

Protein samples were alkylated with 10 mM iodoacetamide in the dark at 37°C for 30m and subsequently digested with trypsin (Promega, 50 ng) overnight at 37°C. Digested peptide mixtures were then acidified, partially dried down in a SpeedVac (Savant) and desalted using a home-made C18 (3M Empore) stage tip filled with 0.4μl of poros R3 (Applied Biosystems) resin. Bound peptides were eluted with 30-80% acetonitrile in 0.1% Trifluoroacetic acid and partially dried to prepare for LC-MS/MS. Liquid chromatography was performed on a fully automated Ultimate 3000 RSLC nano System (Thermo Scientific) fitted with a 100 μm x 2 cm PepMap100 C18 Nano Trap column and a 75 μm×25 cm reverse phase C18 nano column (Aclaim PepMap, Thermo Scientific). Samples were separated using a binary gradient consisting of buffer A (2% MeCN, 0.1% formic acid) and buffer B (80% MeCN, 0.1% formic acid), with a flow rate of 300 nL/min. The HPLC system was coupled to a Q Exactive Plus mass spectrometer (Thermo Scientific) equipped with a nanospray ion source. The mass spectrometer was operated in standard data dependent mode, performed MS full-scan at 350-1600 m/z range, with a resolution of 70000. This was followed by MS2 acquisitions of the 15 most intense ions with a resolution of 17500 and NCE of 27%. MS target values of 1e6 and MS2 target values of 1e5 were used. Isolation window of precursor was set at 1.5 Da and dynamic exclusion of sequenced peptides was enabled for 30s. The acquired MS/MS raw files were searched using Sequest (Proteome Discoverer v2.1) search engine. MS/MS spectra were searched against the *M. tuberculosis* EccA1 sequence including a contaminant database. A list of EccA1 peptides were selected from the result for Parallel Reaction Monitoring (PRM) experiment. For PRM, the mass spectrometer performed a MS full-scan at 400-1600 m/z range, with a resolution of 35000. This was followed by one MS2 acquisitions with a resolution of 35000. MS target values of 3e6 and MS2 target values of 5e5 were used. Isolation window of precursor was set at 0.7 Da.

### Generation of plasmids and mutants

Primers and plasmids are listed in Supplementary Tables 1 and 3. *M. tuberculosis* EccA1 was amplified from the pRD1-2F9 cosmid (Kind gift from Roland Brosch, Institut Pasteur), and cloned into the pH3c-LIC backbone (PSI:Biology-Materials Repository). EccA1 C204S and C204V point mutants were generated via site directed mutagenesis of the resulting pH3c-LIC EccA1 plasmid. To generate the pMOFXh complementation plasmid, a dsDNA fragment was designed with *ccdB* and *cam* genes flanked by SapI restriction sites and regions complementary to the 30 bp surrounding the HpaI and EcoRV sites in pMV306hsp. This fragment was then cloned into pMV306hsp digested with HpaI and EcoRV via *in vivo* Assembly[42]. EccA1, C204S, and C204V complementation constructs were cloned from the pH3c-LIC plasmid into pINIT_kan, and were subsequently flipped into pMOFXh using FX cloning[43].

## Figure Legends

**Figure S1:**
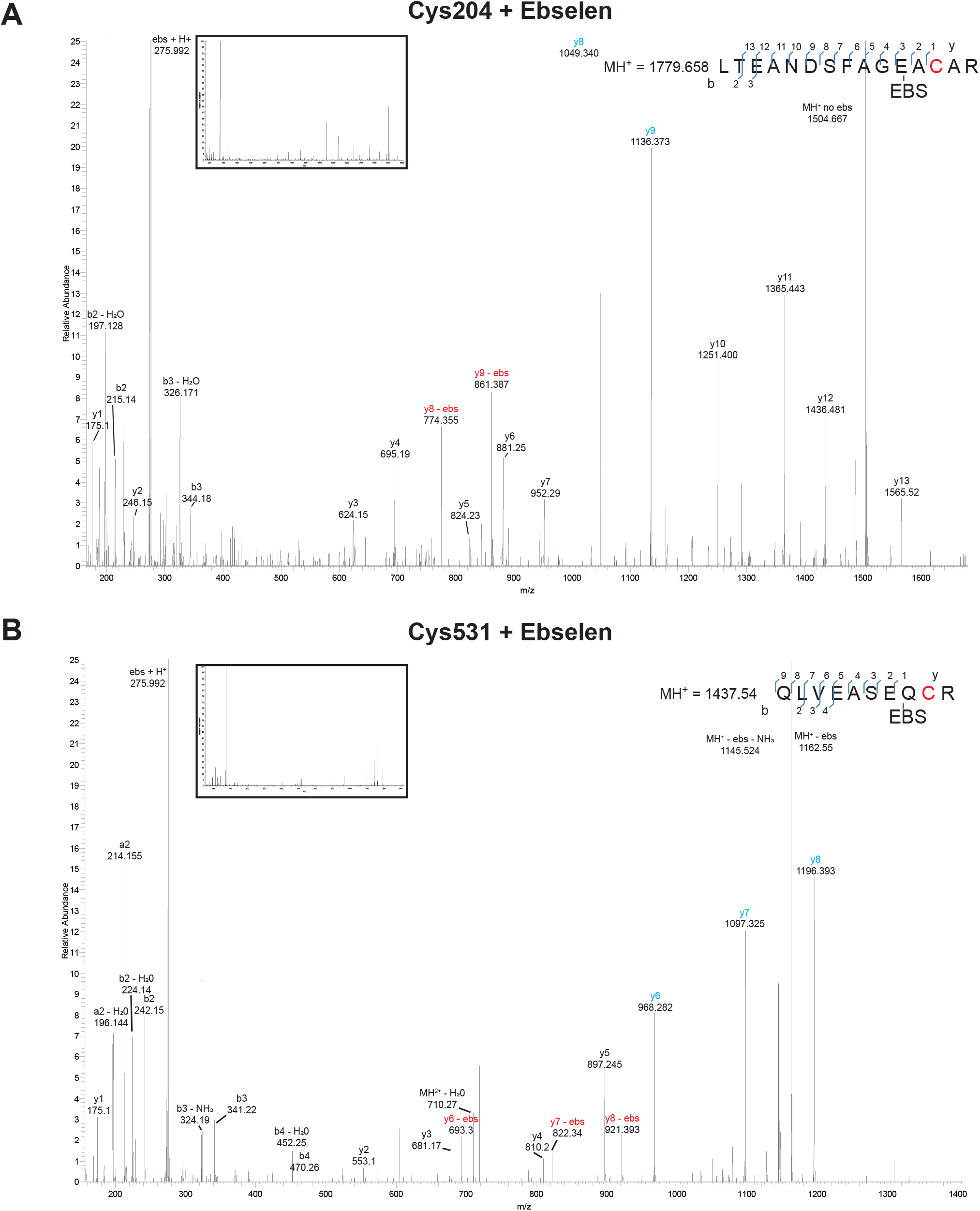
Mass spectrometry reveals peptides confirming ebselen modification of Cys204 and Cys531. **A and B:** MS/MS spectra of ebselen bound Cys204 and Cys531 generated from EccA1 treated with 8 μM ebselen. Fragmentation map is included in the top right corner. Spectra are scaled for readability, unscaled are inset. Masses of different b/y ions are displayed. y-ions include mass of ebselen, unless otherwise stated. Ionized ebselen is visible at 275.992 m/z, as some ebselen-cysteine bonds are broken during peptide fragmentation. As a result, fragments are observed with (cyan) and without (red) bound ebselen, separated by ebselen’s nominal mass (275).

**Figure S2:**
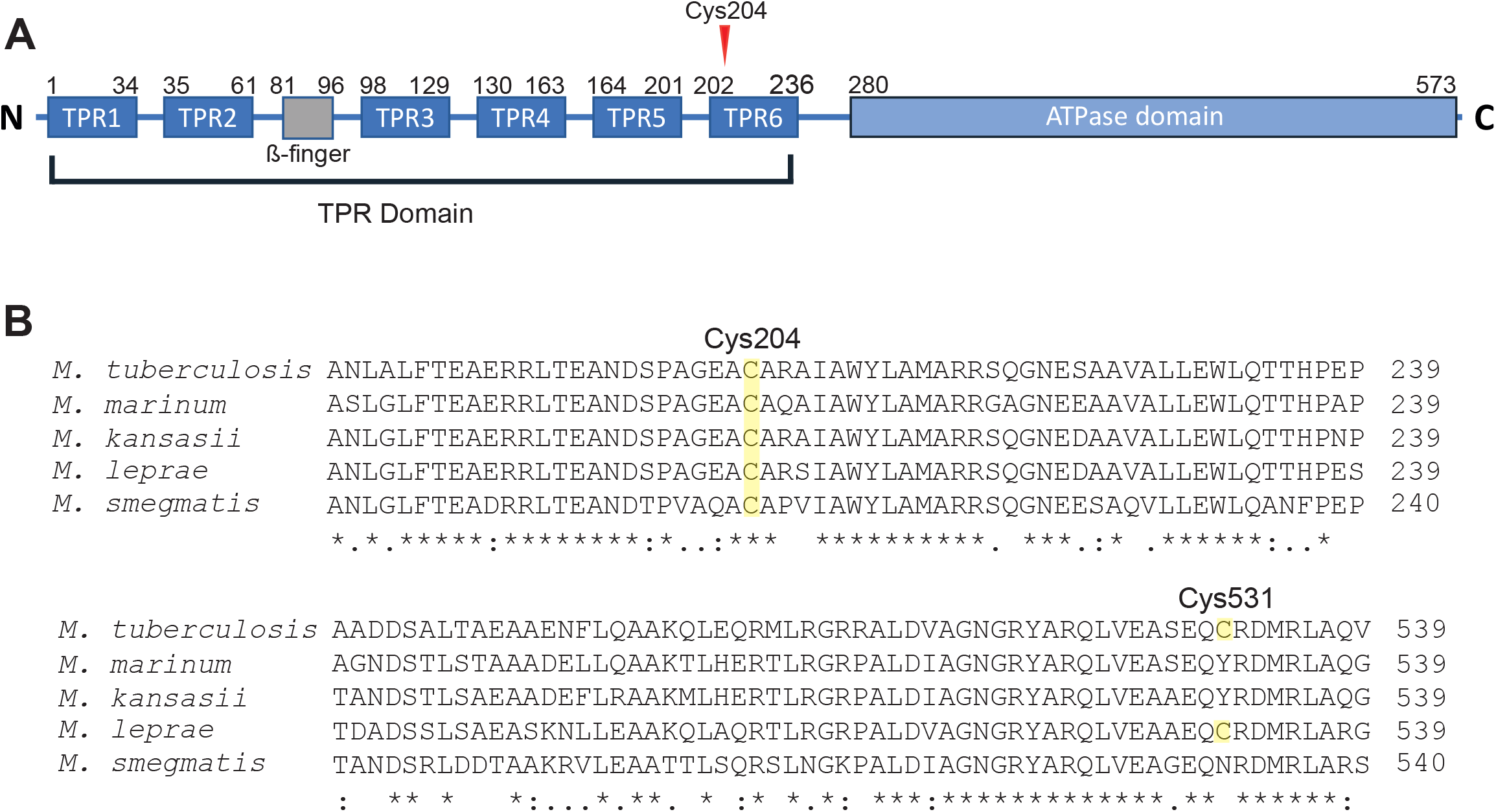
Cys 204 is conserved between *M. tuberculosis* and *M. marinum*. **A:** Diagram of *M. tuberculosis* EccA1’s N-terminal TPR domain including β-finger insertion, and its C-terminal AAA+ ATPase domain. Redrawn from [26].**B:** Clustal Omega multiple sequence alignment of EccA1 homologs with Cys204 and Cys531 annotated. * indicates positions with fully conserved residues. (:)indicates conservation between groups of strongly similar properties (> 0.5 in Gonnet PAM 250 matrix). (.) indicates conservation between groups of weakly similar properties (≤0.5 Gonnet PAM 250 matrix)

