## Supplementary Tables & Figures for "Ebselen attenuates mycobacterial virulence through inhibition of ESX-1 secretion"

**This PDF file includes:**
Supplementary Figures 1 to 2

Supplementary Tables 1 to 3

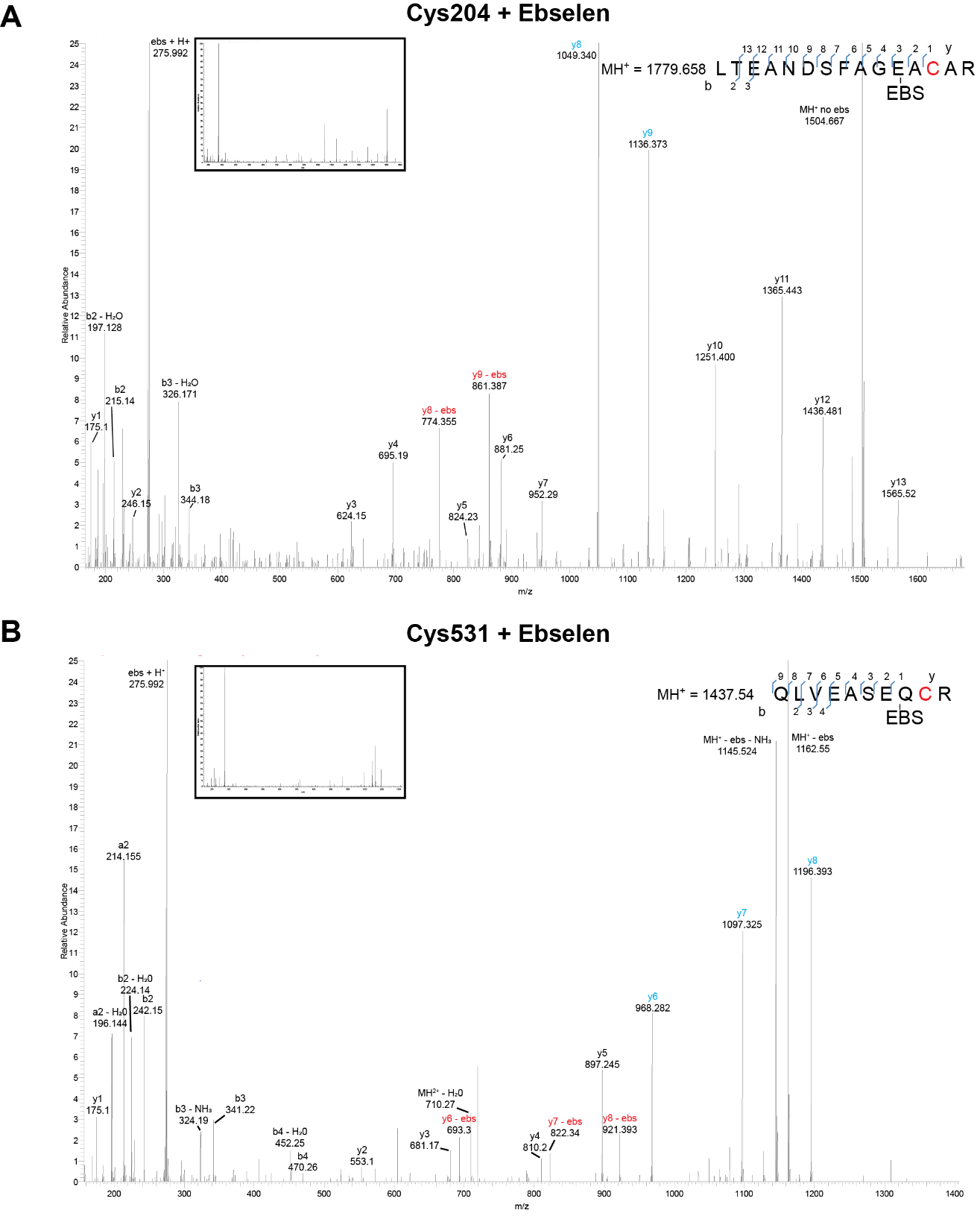

Figure S1: Mass spectrometry reveals peptides confirming ebselen modification of Cys204 and Cys531. **A and B:** MS/MS spectra of ebselen bound Cys204 and Cys531 generated from EccA1 treated with 8 μM ebselen. Fragmentation map is included in the top right corner. Spectra are scaled for readability, unscaled are inset. Masses of different b/y ions are displayed. y-ions include mass of ebselen, unless otherwise stated. Ionized ebselen is visible at 275.992 m/z, as some ebselen-cysteine bonds are broken during peptide fragmentation. As a result, fragments are observed with (cyan) and without (red) bound ebselen, separated by ebselen’s nominal mass (275).

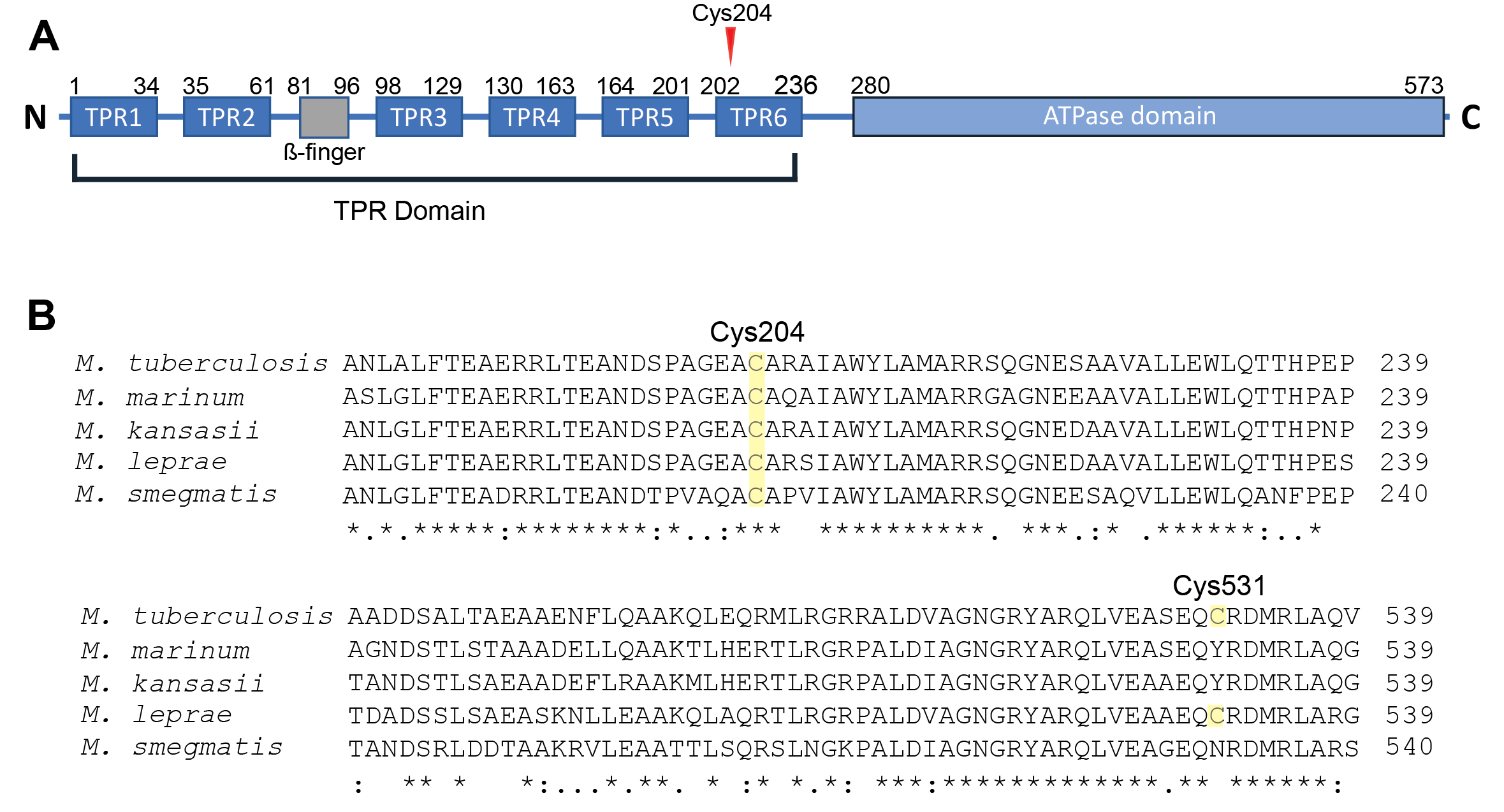

Figure S2: Cys 204 is conserved between *M. tuberculosis* and *M. marinum*. **A:** Diagram of *M. tuberculosis* EccA1’s N-terminal TPR domain including β-finger insertion, and its C-terminal AAA+ ATPase domain. Redrawn from (1)*.***B:** Clustal Omega multiple sequence alignment of EccA1 homologs with Cys204 and Cys531 annotated*.* * indicates positions with fully conserved residues. (:)indicates conservation between groups of strongly similar properties (> 0.5 in Gonnet PAM 250 matrix). (.) indicates conservation between groups of weakly similar properties (≤0.5 Gonnet PAM 250 matrix)

**Table 1: Plasmids used in this manuscript**

| # | Plasmid | Description | Resistance | Ref. |
| --- | --- | --- | --- | --- |
| 1 | pRD1-2F9 | pYUB412 integrating cosmid containing *M. tuberculosis* ESX-1 locus. | Hygromycin | (2) |
| 2 | pH3c-LIC | Expression vector with a N-terminal 8x His tag with a 3C protease cleavage site for use with ligation independent cloning (LIC). | Kanamycin | (3) |
| 4 | pH3c-LIC EccA1 | Expression vector for purification of *M. tuberculosis* EccA1. | Kanamycin | This Study |
| 5 | pH3c-LIC EccA1 C204V | Expression vector for purification of *M. tuberculosis* EccA1 C204V point mutant. | Kanamycin | This Study |
| 6 | pH3c-LIC EccA1 C204S | Expression vector for purification of *M. tuberculosis* EccA1 C204S point mutant. | Kanamycin | This Study |
| 7 | pMOFXh | Mycobacterial FX expression vector with hsp60 promoter, derived from pMV306_hsp. | Kanamycin | This Study |
| 8 | pMOFXh-EccA1 | pMOFXh containing *M. tuberculosis* EccA1 insert. | Kanamycin | This Study |
| 9 | pMOFXh-EccA1 C204V | pMOFXh expressing *M. tuberculosis* EccA1 C204V insert | Kanamycin | This Study |
| 10 | pMOFXh-EccA1 C204S | pMOFXh expressing *M. tuberculosis* EccA1 C204S insert | Kanamycin | This Study |
| 11 | pTEC31 | Mycobacterial plasmid containing the gene for the fluorescent protein tdTomato under the constitutive  mycobacterial promoter msp12. | Kanamycin | (4) |

**Table 2: Mycobacterial strains used in this manuscript**

| # | Strain | Description | Selection | Fig. | Ref. |
| --- | --- | --- | --- | --- | --- |
| 1 | M strain | Wildtype *M. marinum* | None | 1 | (5) |
| 2 | mc^2^6206 | *M. tuberculosis* H37Rv containing deletions of the essential *leuD* and *panCD* loci. | None | 5 | (6) |
| 3 | *eccA1*::Tn | Transposon mutant 19729 containing transposon disrupting the *eccA1* gene. | Hygromycin | 3 | This study |
| 4 | Δ*eccA1* | *M. marinum* *eccA1*::Tn mutant with the hygromycin cassette removed. | None | 3 | This study |
| 5 | ΔeccA1::EccA1_Mtb_ | ΔeccA1 Mm complemented with pMOFXh-eccA1 | Kanamycin | 3 | This study |
| 6 | Δ*eccA1*::C204V | ΔeccA1 Mm complemented with pMOFXh-eccA1-C204V | Kanamycin | 3 | This study |
| 7 | Δ*eccA1*::C204S | ΔeccA1 Mm complemented with pMOFXh-eccA1-C204S | Kanamycin | 3 | This study |
| 8 | Mm-tdTomato | Strain 1 with plasmid 11. | Kanamycin | 4 | (4) |
| 9 | Mm-ΔRD1-tdTomato | Strain 1 with a deletion syntenic to the RD1 locus in M. tuberculosis transformed with plasmid 11. | Kanamycin | 4 | (7) |
| 10 | mc^2^6206-tdTomato | Strain 2 transformed with plasmid 11. | Kanamycin | 5 | This study |

**Table 3: Primers used in this manuscript**

| Name | Sequence | Use |
| --- | --- | --- |
| WC283 | CAAGGACCGAGCAGCCCCACTGATCGCTTGGCCAGT | Amplification of *M. tuberculosis* EccA1 to generate pH3c-LIC EccA1 |
| WC288 | CCACGGGGAACCAACCCTTATCATTCTCTCAGTTGAGGTGTG | Amplification of *M. tuberculosis* EccA1 to generate pH3c-LIC EccA1 |
| M222 | CCGGTGAGGCGGTTGCGCGCGCCATC | Mutagenesis of pH3C-LIC EccA1 to produce pH3c-LIC EccA1 C204V |
| M223 | GATGGCGCGCGCCGACGCCTCACCGG | Mutagenesis of pH3C-LIC EccA1 to produce pH3c-LIC EccA1 C204V |
| MP094 | CCGGTGAGGCGTCTGCGCGCGCCATC | Mutagenesis of pH3C-LIC EccA1 to produce pH3c-LIC EccA1 C204S |
| MP095 | CCGGTGAGGCGGTTGCGCGCGCCATC | Mutagenesis of pH3C-LIC EccA1 to produce pH3c-LIC EccA1 C204S |
